# Genome-wide scale analyses identify novel BMI genotype-environment interactions using a conditional false discovery rate

**DOI:** 10.1101/2020.01.22.908038

**Authors:** R. Moore, L. Georgatou-Politou, J. Liley, O. Stegle, I. Barroso

**Author notes:** Correspondence should be addressed to Oliver Stegle and Inês Barroso.

## Abstract

Genotype-environment interaction (G×E) studies typically focus on variants with previously known marginal associations. While such two-step filtering greatly reduces the multiple testing burden, it can miss loci with pronounced G×E effects, which tend to have weaker marginal associations. To test for G×E effects on a genome-wide scale whilst leveraging information from marginal associations in a flexible manner, we combine the conditional false discovery rate with interaction test results obtained from StructLMM. After validating our approach, we applied this strategy to UK Biobank (UKBB) data to probe for G×E effects on BMI. Using 126,077 UKBB individuals for discovery, we identified known (*FTO, MC4R, SEC16B*) and novel G×E signals, many of which replicated (*FAM150B*/*ALKAL2,TMEM18, EFR3B, ZNF596-FAM87A, LIN7C-BDNF, FAIM2, UNC79, LAT)* in an independent subset of UKBB (n=126,076). Finally, when analysing the full UKBB cohort, we identified 140 candidate loci with G×E effects, highlighting the advantages of our approach.

---

Genotype-environment interaction (G×E) studies aim to identify variants with trait-effects that vary under different environmental exposures, which can provide biological insights for understanding trait onset and progression. Additionally, such interactions yield modifiable risk factors, which may be easier to target for disease prevention, and G×E analyses help to identify subgroups of the population at greatest risk of disease who would most benefit from lifestyle or drug intervention for disease prevention^1–3^.

Whilst G×E effects have already been identified for different traits, including body mass index (BMI) and obesity^3–15^, the number of robustly detected G×E effects remains small^2^.

G×E scans are not currently conducted on a genome-wide scale due to the prohibitive multiple testing burden that they incur, along with lower power to detect interaction effects compared to marginal association effects^16,17^. Instead, G×E studies are typically based on a two-stage design, where in the first stage variants with marginal association effects are selected and in the second stage, these selected variants are tested for G×E effects. Whilst such independent filtering^18^ reduces the burden of multiple testing, this approach assumes that only variants with sufficiently strong marginal association signals can be subject to interaction effects. Consequently, variants with the strongest and hence most interesting G×E effects may be missed if they fall below the chosen selection threshold for the marginal association signal. This may be particularly problematic if the marginal association signals used are obtained from meta-analyses that combine data across multiple cohorts (in particular for different ancestry groups), which are exposed to different environments. Consequently, there is a need for more flexible approaches that leverage the marginal association signal without the need to perform rigid filtering.

## Results

Here, we combine the conditional false discovery rate (cFDR) with G×E interaction testing using StructLMM^10^ to address the aforementioned limitations of G×E studies. The cFDR has traditionally been applied to leverage the marginal association signal for one trait to improve power in detecting association for a second related trait.

Briefly, the key concept of the cFDR is that an informative covariate allows for estimating and accounting for variations in the distribution of null to non-null variants across all tests^19–21^. The cFDR for a given hypothesis can also be defined as the posterior probability of being null given thresholds on both the P value of the tested hypothesis and the corresponding covariate value^19^.

In this work, we adapt the cFDR to perform genome-wide G×E analysis using the StructLMM interaction test (StructLMM-int)^10^. In this setting, the cFDR allows for leveraging the marginal association signal for the discovery of G×E effects, by performing data driven false discovery. This approach improves the power for G×E analyses, enabling such scans to be conducted at a genome-wide scale by leveraging the information that is contained in the corresponding marginal association signals; however, without the need to select arbitrary filtering thresholds. Intuitively, this cFDR-based approach can be interpreted as a ‘soft’ thresholding equivalent of the commonly employed two-step filtering approach without the need to define rigid thresholds.

### Simulated data

To validate and assess this approach, we initially considered simulated data using 5,000 individuals of European ancestry based on genotypes from the 1000 Genomes Project^22^. Following Moore *et al.*^10^, we simulated interaction effects based on 60 environmental covariates derived from UKBB. We generated phenotypes with 550 independent genetic effects, with varying fractions of phenotypic variance explained by G×E versus marginal effects (*ρ*, **Methods**). For non-zero values of *ρ* the marginal association signal acts as an informative covariate in assigning the probability that a variant has non-null G×E (**Supp. Fig. 1a**). Conversely, under the null, when simulating only marginal genetic effects (*ρ* = 0, **Supp. Fig. 1b**, **Methods**) or when simulating no causal genetic effects (**Supp. Fig. 1c**, **Methods**), the marginal association test results do not act as an informative covariate in assigning the probability that a variant has non-null G×E.

We then assessed the power of the cFDR approach to identify variants with true simulated G×E effects (*ρ* > 0), as well as the power to detect variants within different G×E bins, stratifying based on the strength of the simulated G×E effects (*ρ*). We compared the results to those obtained from conventional FDR (Benjamini-Hochberg^23^), as well as two step-filtering for different selection thresholds (**Methods**). We found that the cFDR approach was better powered than either of these approaches (*ρ* > 0, 1% FDR, **Fig. 1a**); all methods empirically control the FDR (**Fig. 1b**, **Methods**). As expected, two-step filtering performs well for identifying variants with weak G×E effects (*ρ* < 0.5, **Fig. 1a**), whereas the conventional FDR when considering all variants is only competitive for detecting variants with very strong G×E effects (*ρ* = 1, **Fig. 1a**), when the marginal association signal is not informative. In contrast, the cFDR approach performs well in both regimes and in particular overcomes the need to choose a suitable selection threshold for the association signal (conventional FDR corresponds to the 100% threshold), which will vary depending on the (*a priori* unknown) extent of G×E.

**Figure 1.**
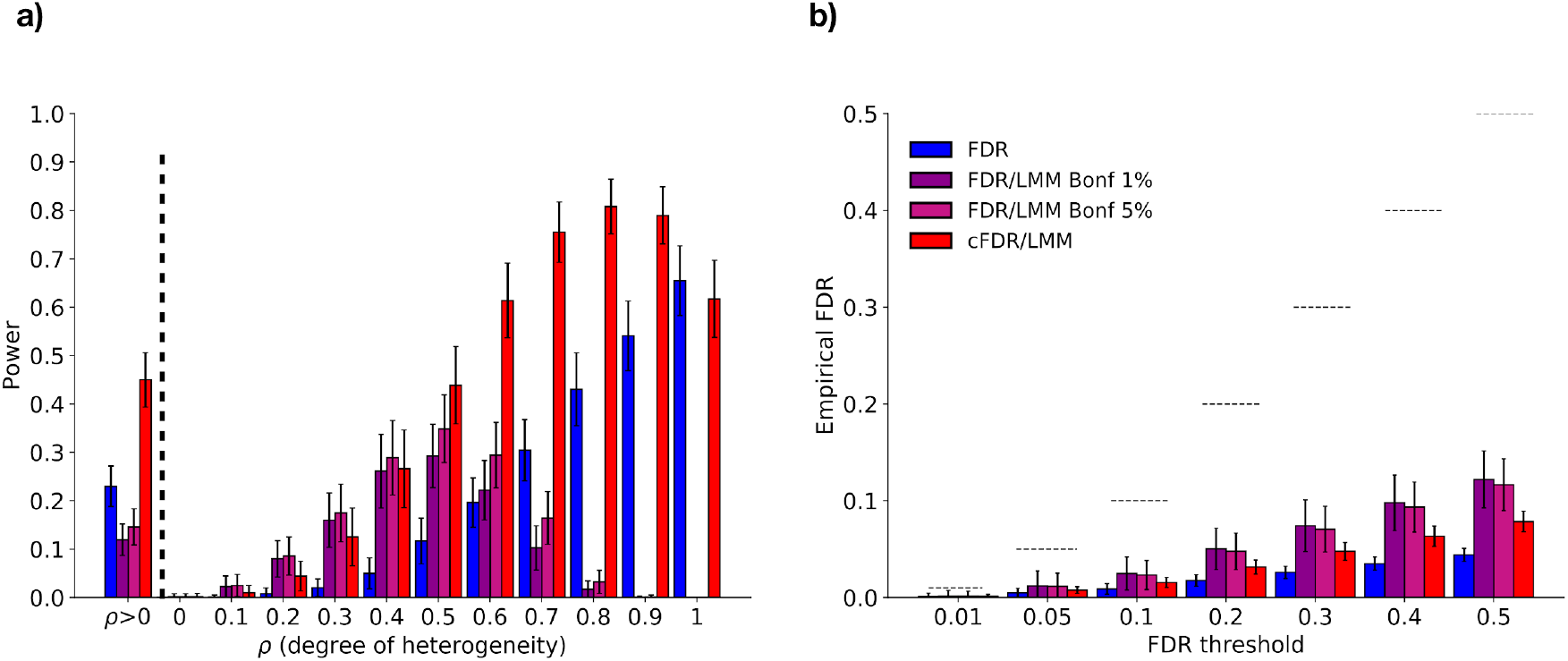
Assessment of power and empirical FDR control of alternative methods using simulated data. (**a**) Comparison of power for detecting simulated interaction effects, considering varying fractions of genetic variance explained by G×E (*ρ*), based on a synthetic European population of 5,000 individuals derived from 1000 Genomes^22^ genotypes (1,650,000 total variants, 550 with non-zero genetic effects; **Methods**). Shown is power to detect variants with G×E effects (FDR < 1%), considering either all variants with G×E (*ρ* > 0, left), or variants stratified by the extent of simulated G×E (right). (**b**) Empirical assessment of the FDR control of alternative methods for increasing thresholds; dashed lines indicate the selected FDR thresholds (**Methods**). Compared were cFDR considering all 1,650,000 tested variants, FDR (Benjamini-Hochberg^23^) considering all 1,650,000 variants and FDR (Benjamini-Hochberg^23^) applied to the subset of variants selected at 1% (3,275 - 5,000 tested variants per repeat experiment) and 5% (3,937 - 5,947 tested variants per repeat experiment) FWER based on marginal association effects, using p-values obtained from the StructLMM interaction test and LMM association test for all three methods. Displayed are the average results across 100 repeat experiments, with error bars denoting plus or minus one standard deviation.

### Application to data from UK Biobank

Next, we applied the cFDR approach to UKBB to identify genetic variants with G×E effects on BMI. We considered 252,153 unrelated UKBB individuals of European ancestry for which BMI and 64 lifestyle environmental factors (diet-related factors, three factors linked to physical activity and six lifestyle factors, modelled as gender and age adjusted, similar to Moore *et al.*^10^; **Methods**), were available in the full release of UKBB. We split the cohort randomly into a discovery and validation set consisting of 126,077 and 126,076 individuals, respectively and considered 7,515,856 low-frequency and common variants (imputed variants, MAF > 1%, **Methods**) for analysis.

On the discovery dataset, we first applied a genome-wide G×E scan using conventional FDR, which identified 78 variants mapping to three loci (*ACTBL2-PLK2, FTO, SDCCAG3P1-MC4R*) with G×E effects (Benjamini-Hochberg; FDR < 5%, **Methods**; **Fig. 2a**, **Supp. Fig. 5a**, **6a**, **7**, **Supp. Table 1**).

**Figure 2.**
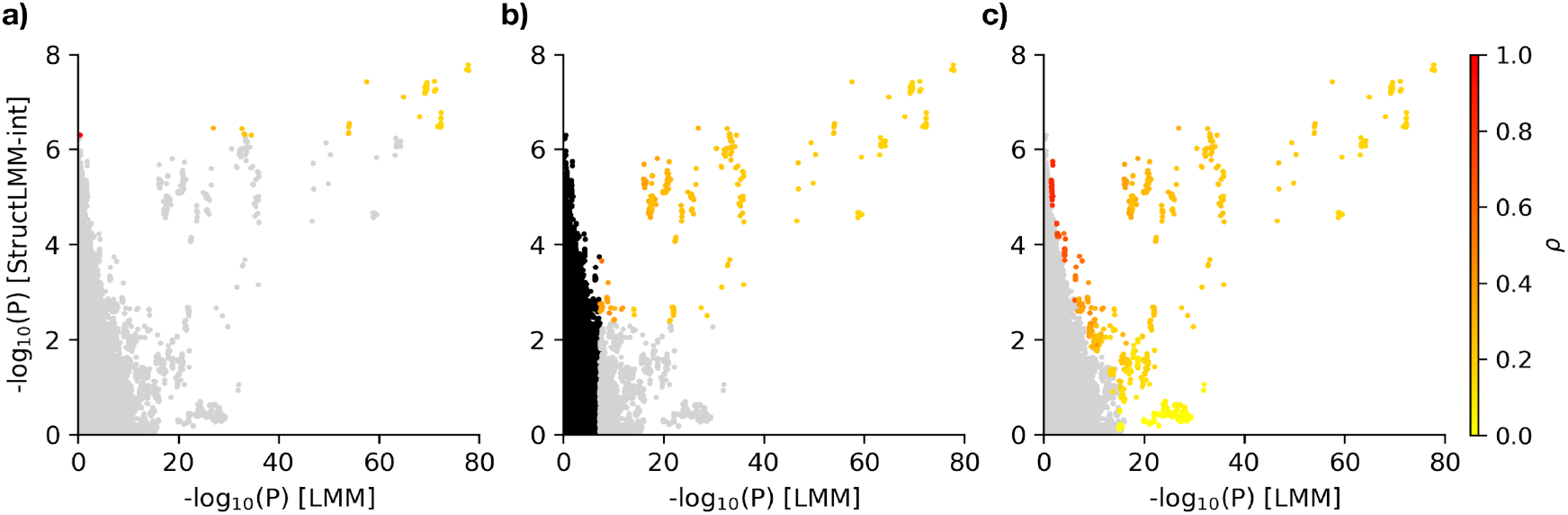
Comparison of variants with G×E effects in the discovery set of UKBB identified by alternative methods. Scatter plots of negative log StructLMM interaction P values (y-axis) versus negative log LMM association P values (x-axis, 7,515,856 variants), obtained from UKBB BMI data using the discovery set of individuals (n = 126,077). Variants with G×E effects (FDR < 5%) are highlighted, with colour denoting the estimated fraction of the genetic variance due to G×E (*ρ*). Considered were (**a**) genome-wide analysis where all 7,515,856 variants were tested for interaction effects (78 variants mapping to three loci < 5% FDR; Benjamini-Hochberg, **Methods**, **Supp. Table 1**), (**b**) two-step filtering considering variants with genome-wide significant marginal association signals (P < 5×10^−8^; 3,767 variants) for interaction testing (variants in black not tested, 330 variants mapping to eight loci < 5% FDR; Benjamini-Hochberg **Methods**, **Supp. Table 1**) and (**c**) a genome-wide analysis of all 7,515,856 variants using cFDR (964 variants mapping to 29 loci < 5% cFDR, **Methods**, **Supp. Table 1**). Loci were defined using LD clumping (r^2^ < 0.1, +/− 500kb; **Methods**).

Next, we applied a two-step filtering, whereby 3,767 variants with genome-wide significant BMI association effects (P < 5×10^−8^, LMM, **Supp. Fig. 4**) were considered for interaction testing. This identified 330 variants corresponding to eight distinct loci, including two (near *FTO* and *SDCCAG3P1-MC4R)* of the three loci identified above, with significant G×E effects (Benjamini-Hochberg; FDR < 5%, distinct based on distance clumping of +/−500kb and LD r^2^ > 0.1, **Methods**; **Fig. 2b**, **Supp. Fig. 5b**, **6b**, **8**, **Supp. Table 1**). Notably, because of the assumption that tested variants have substantial marginal effects, by design this approach only identified variants with weak to moderate G×E effects (*ρ* < 0.556 for all 330 variants with significant G×E effects, Benjamini-Hochberg FDR < 5%).

Finally, we applied cFDR, conditioning the genome-wide G×E test statistics on the corresponding BMI marginal association signal, noting that the marginal association signal is indeed an informative covariate (**Supp. Fig. 3**). This yielded 964 variants corresponding to 29 loci with significant G×E effects (cFDR < 5%, +/−500kb, r^2^ > 0.1, **Methods**; **Fig. 2c**, **Supp. Fig. 5c**, **6c**, **9**, **Supp. Table 1**). As well as identifying all loci (and their tag variants) found by the two-step-filtering approach, the cFDR approach identified 21 additional loci, many of which have moderate to strong G×E effects (**Fig. 2c**, **Supp. Fig. 9**, **Supp. Table 1**) with weak to no marginal BMI association effects (**Supp. Fig. 5c**, **Supp. Table 1**), clearly highlighting the potential gains of using the cFDR approach.

To further assess the additional associations identified, we assessed the replication of the variants detected by the cFDR approach for replication in the independent validation set of individuals (n = 126,076), again using the StructLMM interaction test.

Globally, we observed that variants with G×E identified in the discovery set were enriched for small p-values in the validation set, which includes the 21 loci that were exclusively identified by the cFDR (**Fig. 3a**, **Supp. Fig. 10**). Consistent with this, the tail strength^24^ (deviation measure between the observed and expected set of p-values) for the 29 cFDR loci exceeded the chance expectation (empirical P < 0.002 based on 500 matched sets of null variants^25^, **Methods;** P < 0.002 for the 21 loci), which is similar to loci identified by the two-step filtering approach (P < 0.002).

**Figure 3.**
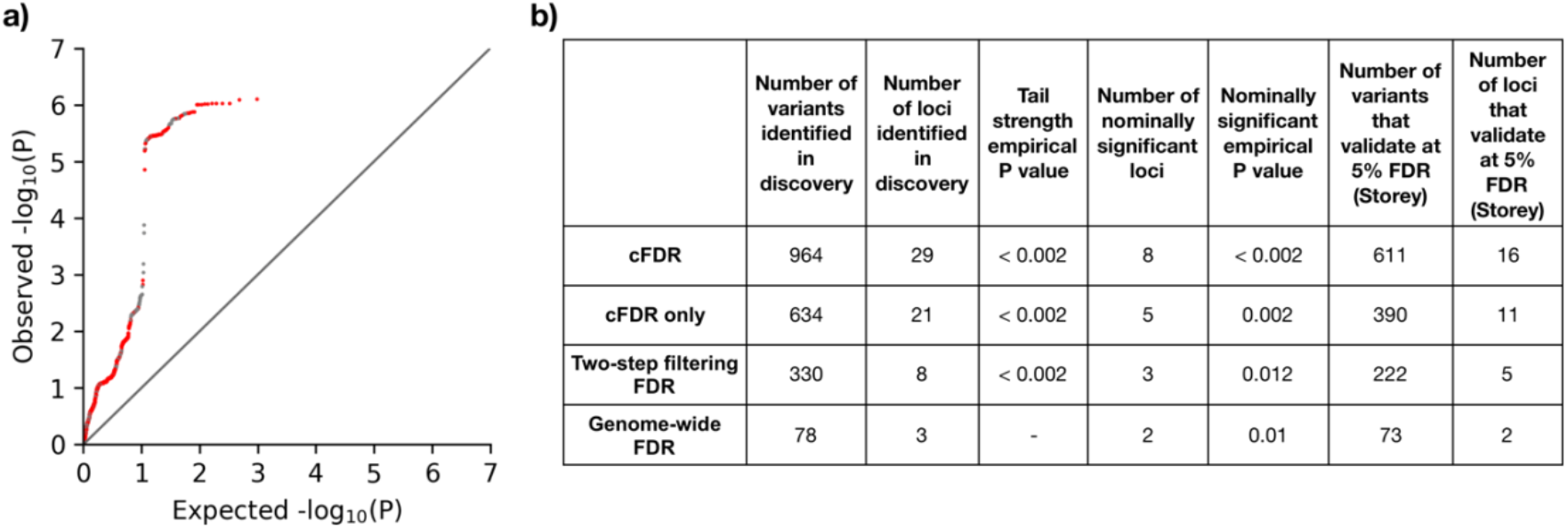
Replication of variants with GxE in the validation datasets. (**a**) QQ plot of negative log P values from the StructLMM interaction test evaluated in the validation dataset (n = 126,076) for the 964 variants identified by the cFDR approach in the discovery analysis. The 330 variants also identified by the genome-wide FDR and/or the two-step filtering in the discovery set are displayed in grey whilst the 634 variants exclusively identified by the cFDR approach are displayed in red. (**b**) Table summarising the validation metrics considered. Columns 1 and 2 state the number of variants and loci (LD clumped loci, r^2^ < 0.1 within +/−500kb), respectively with significant G×E effects (FDR 5%) identified in the discovery set of UKBB individuals (n = 126,077); column 3 displays the significance of the tail strength of the replication P values (based on 500 sets of matched null variants; **Methods**); column 4 states the number of loci identified in discovery that are nominally significant (StructLMM interaction P < 0.05) in validation set, column 5 the corresponding empirical P value based on 500 sets of matched variants (**Methods**), columns 6 and 7 the number of variants that validate the number of variants and loci respectively that validate (Storey FDR 5%^26,27^). ‘cFDR only’ results are based on variants/loci that were only found using the cFDR multiple testing correction.

In addition, the lead variant is nominally significant (P < 0.05) in the validation dataset for eight (*SEC16B*, *ALKAL2-TMEM18*, *ZNF596-FAM87A*, *FAIM2*, *LAT*, *FTO*, *SDCCAG3P1-MC4R and POFUT2)* of the 29 discovered loci, which exceeds the chance expectation (empirical P < 0.002, using 500 matched sets of null variants, **Methods**). In comparison, only three (*FTO*, *SDCCAG3P1-MC4R* and *LAT*) of eight loci for the two-step filtering approach (empirical P = 0.012, using 500 matched sets of null variants, **Methods**) and two (*FTO*, *SDCCAG3P1-MC4R)* of three loci for the genome-wide FDR approach (empirical P = 0.01, using 500 matched sets of null variants, **Methods**) are nominally significant (P < 0.05) in the validation dataset. Considering the 21 loci unique to the cFDR approach in discovery, the lead variant is nominally significant (P < 0.05) for five (*SEC16B, ALKAL2-TMEM18, ZNF596-FAM87A, FAIM2 and POFUT2)* of the 21 loci (empirical P = 0.002, using 500 sets of matched variants, **Methods**). These results indicate that the rate of replication of variants discovered using the cFDR approach is as good as using a conventional two-step filtering pipeline.

Even higher numbers of loci replicated when using Storey’s Q values (estimates the fraction of non-null variants and does not assume it is 1 as for the Benjamini-Hochberg approach)^26,27^, providing further support for using the cFDR when testing for G×E. Sixteen of the loci identified with the cFDR approach replicate (FDR < 5%), whilst in comparison, only five and two of the loci identified by the conventional filtering approach and the genome-wide FDR approach replicate (both are a subset of those that replicate based on the cFDR approach), such that ten loci (*SEC16B*, *ALKAL2*-*TMEM18* two distinct signals, *ZNF596-FAM87A, SEM4AD-GADD456, ITGB1-NRP1, FAIM2, SDCCAG3P1-MC4R two distinct signals, POFUT2)* are identified and replicate solely through the use of the cFDR (**Supp. Table 1**). *BDNF* is identified both by cFDR and the two-step filtering approach but only replicates in the cFDR approach (**Supp. Table 1**).

Finally, having validated the approach, we took advantage of the full set of 252,188 individuals in UKBB, to increase power to identify additional candidate loci with G×E effects on BMI. The cFDR approach identified 140 loci with significant G×E effects (cFDR < 5%, +/−500kb, r^2^ > 0.1, **Methods**, **Supp. Fig. 11 c**, **f**, **Supp. Fig. 13c**, **Supp. Table 2**). In comparison, the genome-wide and conventional two-step filtering approaches identified six and 23 loci with significant G×E effects (FDR < 5%, +/−500kb, r^2^ > 0.1, **Methods**, **Supp. Fig. 11**, **Supp. Fig. 12**, **Supp. Fig. 13**, **Supp. Table 2**), respectively.

Among the 140 loci identified from the full UKBB dataset, in addition to loci overlapping those identified in the discovery and validation datasets, several others merit additional validation in other large biobanks. Of particular interest are ten loci where marginal association with BMI is weak (P > 10^−3^) but with strong evidence of G×E (P < 10^−5^) and where *ρ* is large (> 0.5) (**Supp. Table 2**). Among these is *CADM2*, a locus previously associated with BMI, obesity and adiposity traits^28–32^, in addition to reported associations with physical activity^33^, risk taking behaviour, alcohol, smoking and many other behavioural and cognitive traits^34^, it has recently been suggested to form a link between psychological traits and obesity^35^.

## Discussion

To date, G×E discovery efforts have lagged behind genetic association studies due to limitations in available methods, and sample sizes. Genome-wide G×E scans have a high multiple-testing cost and are therefore limited in power. In contrast, two-step filtering approaches whereby only genetic variants with prior evidence of marginal association with a given trait or disease of interest are tested for G×E effect, limit the multiple-testing burden but are constrained by the number of variants tested and the need to set arbitrary association significance thresholds for filtering. To overcome these limitations, we describe the use of the conditional false-discovery (cFDR) approach, aligned with our recently described StructLMM interaction test method^10^, to perform a genome-wide search for G×E effects whilst avoiding the high multiple testing burden.

Using both simulated data and UKBB data to identify novel loci with evidence of G×E effects on BMI, we highlight two significant advantages of the cFDR approach compared to genome-wide FDR, or two-step filtering approaches. Firstly, cFDR allows a genome-wide G×E exploration whilst retaining greater discovery power compared to conventional genome-wide FDR. Secondly, variants found uniquely by cFDR validate, in an independent dataset, at an equivalent rate to those identified by the conventional hard-filtering approach, demonstrating the utility of the approach to discover variants with a previously unsuspected effect on the trait in question. We further demonstrate that focusing on variants with marginal associations may paradoxically limit the ability to discover variants with the highest G×E effects, because many of these will either have been filtered out in large meta-analyses due to large heterogeneity between contributing datasets, or they will have insufficient evidence for trait-association in combined datasets to be taken forward for G×E testing. This is demonstrated by the two-step filtering approach only identifying loci with low *ρ* value, whereas those loci identified by the cFDR approach have a greater distribution of *ρ* values (**Fig. 2**, **Supp. Fig. 11**).

Using the cFDR approach in combination with StructLMM, we detected 16 replicating loci (only five of which were identified and validated with a conventional two stage design) with evidence of G×E effects on BMI. Two (*FTO* and *MC4R*), had been previously established^5,7,9,10,15,36^, and were also detected when applying genome-wide FDR or two-step filtering approaches. In addition, for the first time we demonstrate evidence of G×E effects on BMI, in *EFR3B*, *LINC7C* and *UNC79*, replicating loci overlapping between cFDR and two-step filtering approaches.

Furthermore, eleven replicating loci were uniquely identified by the cFDR approach (**Supp. Table 1**). Amongst these eleven loci, is *SEC16B*, a positive control for which previous secondary analyses provided some evidence for an interaction (P = 0.025) with physical activity in Europeans^36^, and in Hispanics^37^ and identified more recently by us using the StructLMM interaction test with a sample size approximately twice as large, though we note it contains the samples used here^10^. Notable loci with new evidence for G×E effects on BMI include *ALKAL2, TMEM18 and FAIM2. ALKAL2* and *FAIM2* were also detected when using a two-step filtering approach but only when the entire UKBB dataset was used. *ALKAL2* (previously called *FAM150B)* lead variant rs62107263 (MAF= 22%) is in D’=1 with rs62107261 (MAF=4%), previously associated with obesity and body fat distribution^32,38^, lung function^39^ and smoking status^39^. In *TMEM18* the variant with G×E effects (rs74676797) is in perfect LD (r^2^ = 1) with rs13021737 previously shown to associate with BMI and obesity^29,40,41^. Finally, in *FAIM2* (rs7132908), the same variant has been previously associated with childhood obesity^41^, severe obesity^32^, BMI^39^ and alcohol consumption^42^. The latter is interesting as the environment with the most evidence of driving the interaction effect is alcohol frequency in women (**Supp. Fig. 14**).

In summary, we describe a novel approach that uses the cFDR to discover G×E effects genome-wide without incurring the high multiple testing burden. While we describe an application that combined the cFDR approach with our previous StructLMM interaction test^10^, it is important to note that the cFDR approach can be combined with any G×E test and therefore has wide applicability. Additionally, we note that while we considered marginal association results with BMI as our informative covariate, it is equally valid to use marginal association results with one trait to test for G×E interactions on a second trait, for example, we could use marginal associations with BMI, to look for G×E on type 2 diabetes risk.

## Online methods

### Statistical tests

#### LMM

A conventional linear mixed model (LMM) to test for marginal associations can be cast as

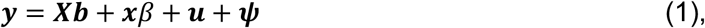

where *β* is the effect size of the focal variant ***x***, ***X*** denotes the fixed-effect design matrix of *K* covariates and ***b*** the corresponding effect sizes. The variable ***u*** denotes additive (confounding) factors, and ***ψ*** denotes i.i.d. noise. The random effect component ***u*** and the noise vector ***ψ*** follow multivariate normal distributions, 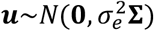 and 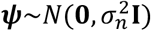, where the covariance matrix **Σ** reflects the covariance of environment across individuals. **Σ** is estimated using a linear covariance function such that **Σ** = **EE**^*T*^, where **E** is an *N* × *L* matrix composed of *L* environmental exposures for each of the *N* individuals (see Moore *et al.*^10^ for full details). Marginal association tests for non-zero effects of the focal variant correspond to alternative hypothesis *β* ≠ 0.

#### StructLMM interaction test

The StructLMM interaction test generalises the LMM by introducing per-individual effect sizes of the focal variant due to G×E and can be cast as

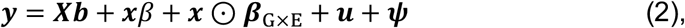

where ⊙ denotes the Hadamard product and ***β***_G×E_ is a per-individual allelic effects vector and follows a multivariate normal distribution with environment covariance 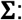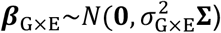, where **Σ** = **EE**^*T*^ as described above. As a result the interaction test, with alternative hypothesis 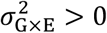, tests for G×E effects due to potentially multiple environmental variables (see Moore *et al.*^10^ for full details). It can be seen that under the null StructLMM reduces to the LMM described above.

#### Fraction of genetic variance explained by G×E (*ρ*)

In this study, 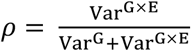 is defined as the fraction of genetic variance that is explained by G×E, where Var^G^ denotes the fraction of the variance explained by marginal genetic effects and Var^G×E^ variance due to G×E. An estimate of *ρ* can be obtained by maximizing the marginal likelihood of Eq. (2) (see Moore *et al.*^10^ for details).

#### The conditional false discovery rate (cFDR)

The cFDR is an estimate of the posterior probability of a primary trait of non-G×E interaction given p-value thresholds for hypothesis tests of G×E interaction and for a secondary trait of phenotype-genotype (G) association. In this work, p-values for the ‘primary trait’ are obtained from application of the StructLMM interaction (StructLMM-int) test (see above) and the conditional informative covariate trait as the association test results, obtained from LMM (see above, Moore *et al.*^10^ description of LMM-Renv for full details).

Formally, denoting 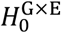 as a null hypothesis of non-G×E, *P* as a random variable corresponding to the p-value from a hypothesis test for G×E from StructLMM, and *Q* as a p-value from a hypothesis test for phenotype association, the cFDR is a function of thresholds *p*, *q*:

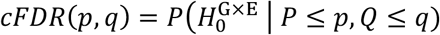

Given observed (*P*, *Q*) values (*p*_1_, *q*_1_), (*p*_2_, *q*_2_), …, (*p*_*n*_, *q*_*n*_) for each SNP, ranking hypothesis by *cFDR*(*p*_1_, *q*_1_), *cFDR*(*p*_2_, *q*_2_), …, *cFDR*(*p*_*n*_, *q*_*n*_) may sort null and non-null SNPs more effectively than ranking by p-values *p*_1_, *p*_2_, …, *p*_*n*_ alone. The best possible metric by which to rank SNPs can be shown^19^ to be:

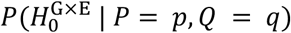

but this quantity is difficult to estimate, whereas the cFDR is readily and partly-consistently estimable from empirical cumulative density function of *P* and *Q*.

The cFDR can be seen to be related to the Benjamini-Hochberg procedure. The Benjamini-Hochberg procedure controlling FDR at *α* is roughly equivalent to rejecting the null for all *i* such that:

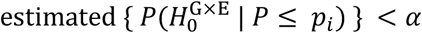

A corresponding rule for the cFDR does not hold^21^; rejecting 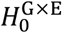 whenever estimated {*cFDR*(*p*, *q*)} < *α* does not control the FDR at *α*.

We control type-1 error rate (as FDR) using an approach proposed by Liley *et al.*^19^ (code available at https://github.com/jamesliley/cfdr, using options mode = 2 and adj = T). Briefly, in this method, type-1 error rate (as FDR) is controlled by using the estimated cFDR to define a map from the unit square (the domain of *P*, *Q*)) to the real line. This enables ‘contours’ of cFDR to be drawn through each p-value pair of interest and the regions within integrated with respect to an estimated distribution of the p-value pairs under 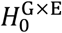, thereby transforming the set of pairs of p-values into a set of single p-values against 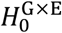 which encompass the additional information from the covariate (and can be used in the Benjamini-Hochberg procedure to control FDR). For correct calibration of the resultant p-value, the map must be independent of the p-value pairs it is used on, so in each case we fitted the map using only p-values from SNPs in linkage equilibrium with the SNP under investigation.

Both the cFDR and two-stage independent filtering^18^ assume independence of *P* and *Q* under 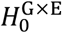 (although both may be adapted if this condition does not hold, and cFDR estimates are reasonably robust to small correlations under the 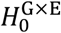^21^). It can be shown that independence of *P* and *Q* holds for nested models (StructLMM interaction test reduces to LMM under the null as described above)^43^. This is in agreement with our simulation experiments that indicate results from the StructLMM interaction test are independent of those from the LMM association test (**Supp. Fig. 1).**

### Simulations

#### Simulation procedure overview

Simulations were based on genotypes of European individuals from the 1000 Genomes project^22^ (phase 1, 1,092 individuals, 379 Europeans), considering 103,527 variants on chromosome 21 (minor allele frequency ≥2%). Following^10,44,45^, 5,000 synthetic genotypes of unrelated individuals were generated for different sample sizes, while preserving the population structure of the seed population (see^46^). We considered 33 environmental exposures using empirical environmental covariates from 70,282 UKBB individuals (based on the Interim release), augmented with gender (binary male indicator vector and binary female indicator vector) and age (continuous vector), resulting in 99 lifestyle covariates and age itself, giving a total of 100 environmental variables. 60 of these environments were selected at random with a subset of 30 environments used to simulate G×E effects and all 60 environments used for testing. These environmental variables were preprocessed as in the UKBB analysis (see below) and randomly assigned to synthetic genotypes (see Moore *et al.*^10^ for full details).

We randomly select 550 segments of approximately 2Mb from chromosome 21, simulating genetic effects from one causal variant. 50 of these segments were simulated with marginal genetic effects only (*ρ* = 0) and G×E effects of varying strengths were simulated for the remaining 500 segments (*ρ* = 0.1, 0.2, 0.3, 0.4, 0.5, 0.6, 0.7, 0.8, 0.9 and 1.0 for 50 segments each). This procedure is repeated 100 times; thereby generating 100 phenotypes, with each the sum of 50 marginal association effects and 500 G×E effects.

#### Conditional QQ plots

QQ plots based on one of the repeat experiments of the StructLMM interaction results were generated for the subset of variants with LMM association p-values < 1 (blue), < 5 ×10^−3^ (orange), < 5 ×10^−5^ (green) and < 5 ×10^−8^ (purple), referred to as conditional QQ plot (**Supp. Fig. 1a**). For comparison, we simulated phenotypes with no dependence between the association and interaction test results, considering i) simulated marginal genetic effects (550 variants with marginal genetic effects but no G×E effects (*ρ* = 0), **Supp. Fig. 1b**) and ii) no simulated genetic effect (**Supp. Fig. 1c**).

#### Assessment of the discovery rate control using simulations

If a variant that lies within a block simulated to have only marginal genetic effects (*ρ* = 0) was declared as having a significant interaction effect, this was defined as a false discovery whilst if a variant lies within a block with simulated causal G×E (*ρ* > 0) was declared as having a significant interaction effect, this was defined as a true discovery. The empirical FDR was then calculated as the number of 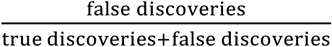, with the average value and standard error over the 100 repeat experiments calculated (**Fig. 1b**).

#### Power simulations

We first calculated power to detect any G×E effects (*ρ* > 0) and then power to detect the subset of simulated causal variants for each of the 11 values of *ρ*. Power was assessed at the 1% FDR, considering variants in linkage disequilibrium with the selected true causal variants (r^2^ ≥ 0.8) as true positives, reporting the average power and standard error across the 100 repeat experiments (each experiment has power 0, 0.002, 0.004, …, 0.996, 0.998, 1 when considering power to detect any interaction effect and 0, 0.02, 0.04…, 0.96, 0.98, 1 when considering power to detect variants at each value of *ρ*, **Fig. 1a**).

#### Comparison methods

We compared the cFDR to alternative multiple test strategies. We considered testing all variants for interaction effects at 1% FDR (Benjamini-Hochberg). We also considered two-stage independent filtering approaches^18^, in which variants were selected for interaction testing based on their association results (LMM) at 1% and 5% FWER. For any of the two-stage filtering approaches, a 1% FDR (Benjamini-Hochberg) was then applied to the subset of variants that were tested for interaction effects.

### Analysis of BMI in UK Biobank

This research has been conducted using the full release of the UK Biobank Resource (Application 14069)^47^. The UK Biobank study has approval from the North West Multi-Centre Research Ethics Committee and all participants included at the time of the analyses provided informed consent to UK Biobank.

#### Data Preprocessing

The data was preprocessed as described in Moore *et al.*^10^. Briefly, BMI phenotype data is ‘Instance 0’ of UKBB data field 21001. Individuals with missing BMI data were discarded and BMI log transformed^15,48^. We considered 21 lifestyle covariates as environments, discarding individuals with outlying or missing environmental data (see Supp. Note in Moore *et al.*^10^ for details). We further discarded individuals of non-British ancestry, related individuals and those that had withdrawn consent at the time of analysis. After these QC procedures on the BMI phenotype, genotype and environmental variables, we had a set of 252,153 individuals for analysis. These individuals were randomly split into a discovery set (n=126,077) and a validation set (n=126,076). Principal components for population structure were those computed by Moore *et al.*^10^, using flashPCA version 2.0^49^ using 147,604 variants as indicated by the field ‘in_PCA’ from the released marker QC file.

#### Genotype data

The genetic variants considered were the same as those used in Moore *et al.*^10^. Specifically, 7,515,856 variants imputed with the HRC panel (build GRCh37) with missingness < 5%, MAF > 1%, HWE P > 1 × 10^−6^, and INFO score r^2^ > 0.4 were considered (see Moore *et al.*^10^ for full details).

#### Environmental covariance and covariates

The environmental matrices **E** for the discovery, validation and analysis using all available samples were generated by augmenting the 21 environmental variables described above with gender (binary male indicator vector and binary female indicator vector) and age (continuous vector), resulting in 63 covariates and age itself was also included, giving a total of 64 environmental factors (see Supp. Note in Moore *et al.*^10^ for details). The environmental covariance was then estimated using standardised environmental variables followed by per-individual standardisation (see Supp. Note in Moore *et al.*^10^ for details). In all analyses, a mean vector, genotype chip, gender, age^2^, age^3^, gender × age, gender × age^2^, gender × age^3^ and ten genetic principal components were included as covariates.

#### Calibration of the considered tests

To check that the association and interaction tests were calibrated under the null, we assessed the empirical calibration using permuted genotype variants (89,166 variants) on chromosome 22 (**Supp. Fig. 2**). Genomic control was calculated as 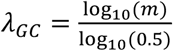; m is the median StructLMM interaction P value.

#### Discovery interaction testing

In these analyses we used only the discovery set of 126,077 individuals.

For the genome-wide FDR approach we applied the 5% Benjamini-Hochberg FDR^23^ to the corresponding G×E results obtained using StructLMM interaction test for all 7,515,856 that passed QC, identifying 78 variants with significant G×E effects.

For the conventional two-step filtering FDR approach, we selected the 3,767 variants with association P < 5×10^−8^ and then applied the 5% Benjamini-Hochberg FDR^23^ to the corresponding G×E results obtained using StructLMM interaction test, identifying 330 variants with significant G×E effects.

For the cFDR approach we applied the 5% cFDR procedure (described above^19^), using 238 folds containing between 30,025 and 44,583 consecutive variants and excluding a fold either side of the focal fold such that the minimum distance between a focal SNP in fold and a SNP out of fold used for the inference was greater than 5,329,663 bp and the maximum LD (based on 100 boundary variants) r^2^ < 9.50×10^−4^ and to the G×E results obtained using StructLMM interaction test for all 7,515,856 that passed QC, using the corresponding association results obtained from LMM for all 7,515,856 variants as the informative covariate, identifying 964 variants with significant G×E effects.

For all variants with significant G×E effects, we estimated *ρ*, which is the fraction of the genetic variance explained by G×E effects (see Moore *et al.*^10^ for details). A value of *ρ* close to 0, means that the effect of variant is largely the same across all individuals with only a small component of the genetic effect dependent on environmental exposures whilst a value close to 1 means that the effect of variant is dependent on the environmental exposures.

LD clumping was performed for each method to identify independent loci by: iteratively (i) selecting the most significant variant (based on the StructLMM interaction results), (ii) removing all variants in LD (r^2^ > 0.1 based on the UKBB discovery set of individuals) within +/− 500kb until no variants were left resulting in three, eight, and 29 clumps (loci) respectively (**Supp. Table 1**). Variants in each loci are visualised in **Supp. Figs. 7**, **8** and **9**, with variants in the clump coloured according to *ρ* and the lead variant is represented by a diamond.

#### Validation

In these analyses we used only the validation set of 126,076 individuals.

We first tested the 964 variants identified with significant G×E effects (cFDR < 5%) in the discovery analysis for interaction effects using the StructLMM interaction test considering the validation set of individuals. QQ plots of the validation StructLMM interaction test results were plotted where (i) the 330 variants also identified by the genome-wide FDR and/or the conventional two-step filtering approach using the discovery set of individuals were coloured in grey (634 variants in red were identified only with the cFDR approach, **Fig. 3**) and (ii) all variants (615 variants) within the eight loci also identified by the genome-wide FDR and/or the conventional two-step filtering approach using the discovery set of individuals were coloured in grey (340 variants in the 21 loci identified only by the cFDR approach are shown in red, **Supp. Fig. 10**).

We assessed the deviation from the null distribution using the tail strength measure^24^ using the lead variant for each locus identified in the discovery analysis based on all 29 cFDR loci, the 8 loci identified by the two-step filtering approach and for the 21 loci identified only by the cFDR approach. Empirical P values were calculated by identifying 500 different matched variants for each lead variant using SNPsnap^25^ (https://data.broadinstitute.org/mpg/snpsnap/match_snps.html), using default settings apart from setting the ‘Number of matched SNPs’ to 4000) and then calculating tail strength for each matched set and subsequently the fraction of the 500 matched sets that are at least as extreme as the observed tail strength measure.

We also counted the number of loci with nominally significant P values (StructLMM interaction test in validation < 0.05) identified by each approach in discovery and also for the 21 loci only identified by the cFDR approach based on lead variants per locus identified in the discovery analysis. Empirical P values were calculated using the same sets of matched variants as were used for the tail strength measure.

Finally, we identified loci that validated for each method at the 5% FDR using Storey’s q value approach^26,27^. This involves estimating the proportion of tested variants that are truly null (*π*_0_) based on the set of variants tested. As the number of variants identified in discovery for the genome-wide FDR and two-step filtering approaches is too low to estimate *π*_0_, the estimate of *π*_0_ obtained for the cFDR approach (0.202, https://www.bioconductor.org/packages/devel/bioc/vignettes/qvalue/inst/doc/qvalue.pdf) was used.

#### Interaction testing using all samples

The data for this analysis was preprocessed as described above but at the time of analysis less individuals had withdrawn consent, resulting in a total of 252,188 individuals. The analysis was conducted analogous to that described in the discovery interaction testing methods section, with the only difference being the larger sample size.

#### Explorative analysis of driving environments at rs7132908

As described in Moore *et al.*^10^, we explored which environments had putative effects on G×E by comparing the log marginal likelihood of the full model to models with individual or sets of environments excluded. We initially assessed the relevance of individual environments based on the log(Bayes factor) of removing single environments (**Supp. Fig. 14a**). To account for correlations between environments, we also used a backwards elimination procedure (see Moore *et al.*^10^ Supp. Note for full details), greedily removing environments until there is evidence that we have selected a full set of environments that can drive the observed G×E effect (**Supp. Fig. 14b**).

## Acknowledgements

This research has been conducted using the UK Biobank Resource (Application Number 14069). R.M. was supported by a PhD fellowship from the Mathematical Genomics and Medicine programme, funded by the Wellcome Trust. IB acknowledges funding from Wellcome (WT206194). J.L. was funded by Johnson and Johnson. O.S. is supported by the BMBF, the European Commission (ERC project DECODE, grant agreement ID: 810296) and the Volkswagen Foundation. L.G.P. was supported by core funding.

## Author contributions

R.M, L.G.P., I.B. & O.S. designed the experiments. R.M. & L.G.P. analysed the simulation data. R.M. analysed the UK Biobank data. J.L. provided code and advice for the cFDR analysis. R.M., L.G.P., I.B. & O.S. interpreted results and wrote the manuscript. All authors read and approved the final version of the manuscript.

