## Supplementary tables and figures for "Genome-wide scale analyses identify novel BMI genotype-environment interactions using a conditional false discovery rate"

**Supplementary Table 1 | Interactions identified by the cFDR, genome-wide FDR and two-step filtering FDR approaches for BMI using the discovery data set (n = 126,077 individuals from UKBB).** Provided as supplementary data file Supplementary_table_1.xlsx

**Supplementary Table 2 | Interactions identified by the cFDR, genome-wide FDR and two-step filtering FDR approaches for BMI using the full set of individuals (n = 252,188 individuals from UKBB).** Provided as supplementary data file Supplementary_table_2.xlsx

### Supplementary figures

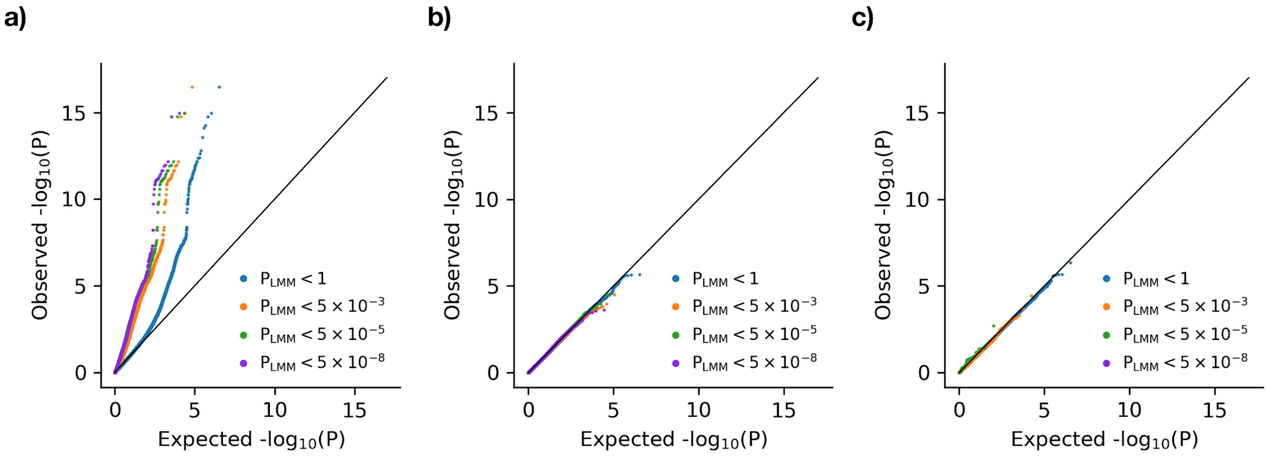


**Supplementary Figure 1 | QQ plots of the StructLMM interaction test conditioned on the LMM association test using simulated data.** QQ plot of negative log P values (1,650,000 variants, 550 causal variants) from the StructLMM interaction test, stratified according to the corresponding association P values obtained from LMM when simulating (**a**) causal variants with varying fractions of phenotypic variance explained by G×E versus marginal effects ($\rho$, **Methods**). In blue are the results for all 1,650,000 variants, in orange for a subset of 33,921 variants with association P values < $5 \times 10^{-3}$, in green for a subset of 11,923 variants with association P values < $5 \times 10^{-5}$ and in purple for a subset of 5,330 variants with association P values < $5 \times 10^{-8}$. Using increasingly stringent association P value thresholds results in a leftward shift of the stratified QQ plots, indicating that association P values are an informative covariate for interaction testing under the alternative, (**b**) only marginal genetic effects ($\rho = 0$), (**c**) no genetic effects. A synthetic European population of 5,000 individuals based on 1000 Genomes Project genotypes was used for all experiments (**Methods**) and the results displayed correspond to one of the 100 experiments conducted to generate **Fig. 1**.

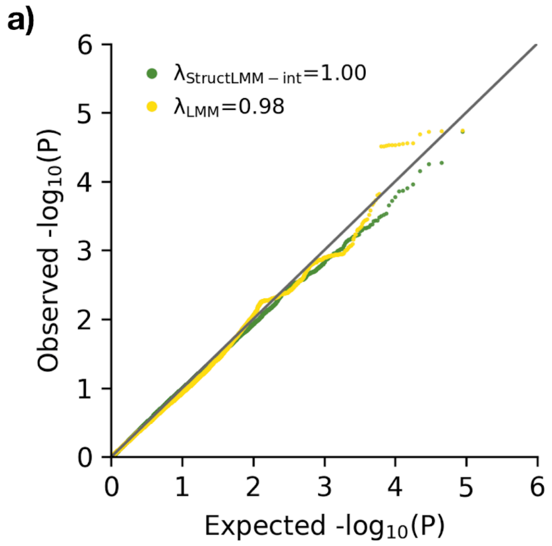


**Supplementary Figure 2 | Calibration of StructLMM interaction and LMM association test on UKBB discovery and validation datasets.** QQ plots of negative log P values based on permuted genetic variants (chromosome 22, 89,166 variants) for (**a**) StructLMM interaction (StructLMM-int, green) and LMM (yellow) tests applied to UKBB BMI phenotype data using the discovery set of individuals (n = 126,077) and (**b**) StructLMM interaction test applied to UKBB BMI phenotype data using the validation set of individuals (n = 126,076). StructLMM interaction and LMM tests were calibrated.

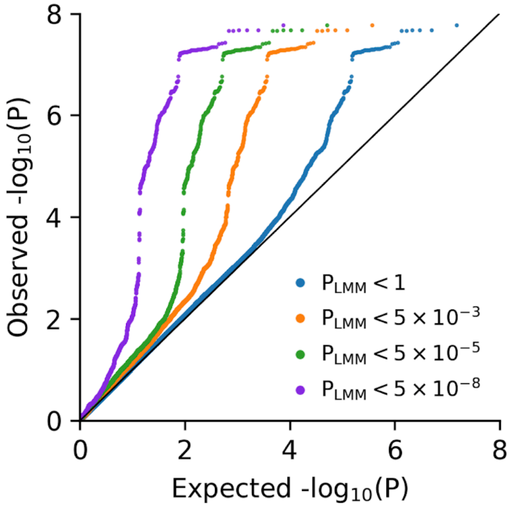


**Supplementary Figure 3 | QQ plot of the StructLMM interaction test conditioned on the association test for UKBB discovery dataset.** QQ plot of genome-wide negative log P values (7,515,856 variants) from the StructLMM interaction test, stratified according to the corresponding association P values obtained using LMM applied to UKBB BMI phenotype data using the discovery set of individuals (n = 126,077). In blue are the results from using all 7,515,856 variants, in orange using a subset of 183,534 variants with association P values $< 5 \times 10^{-3}$, in green using a subset of 25,843 variants with association P values $< 5 \times 10^{-5}$ and in purple using a subset of 3,767 variants with association P values $< 5 \times 10^{-8}$. Using increasingly stringent association P value thresholds results in a leftward shift of the stratified QQ plots, indicating that association P values are an informative covariate for interaction testing under the alternative.

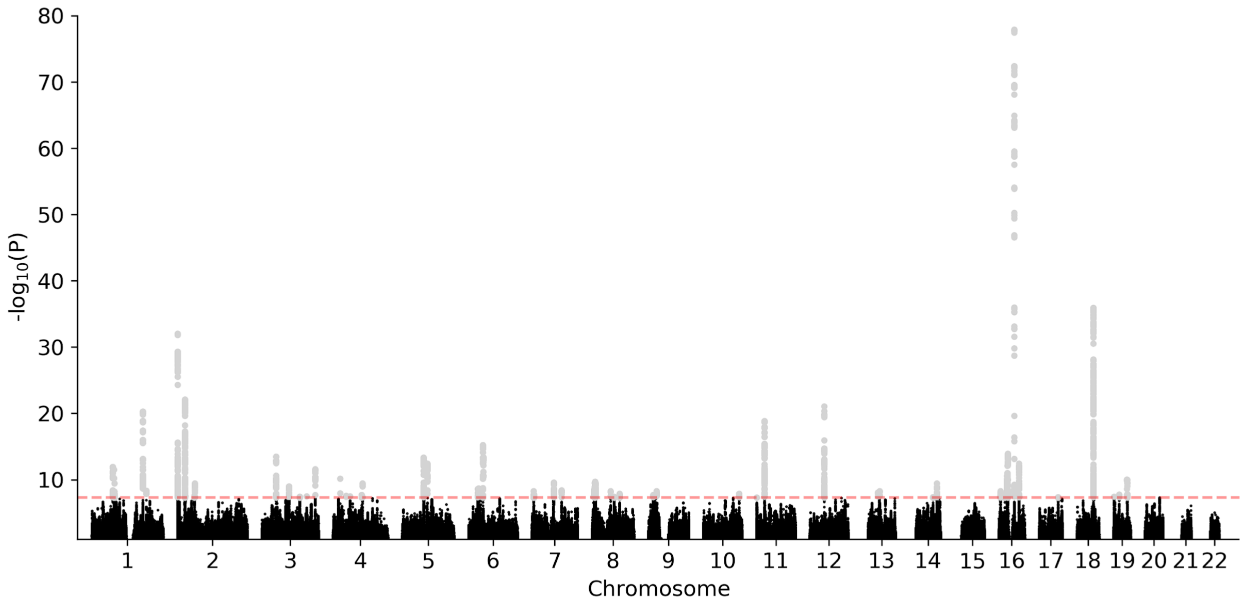


**Supplementary Figure 4 | Association test results for the UKBB discovery dataset.** Manhattan plot showing genome-wide negative log P values (7,515,856 variants) obtained from the LMM association test, applied to UKBB BMI data on the discovery set of individuals (n = 126,077). The dashed red line denotes the genome-wide significance threshold ($P < 5 \times 10^{-8}$). Genome-wide significant variants (grey) are considered for interaction testing by the two-step filtering FDR approach. 3,767 variants, corresponding to 81 loci (LD clumped loci, $r^2 < 0.1$ within +/-500kb, **Methods**) are genome-wide significant.

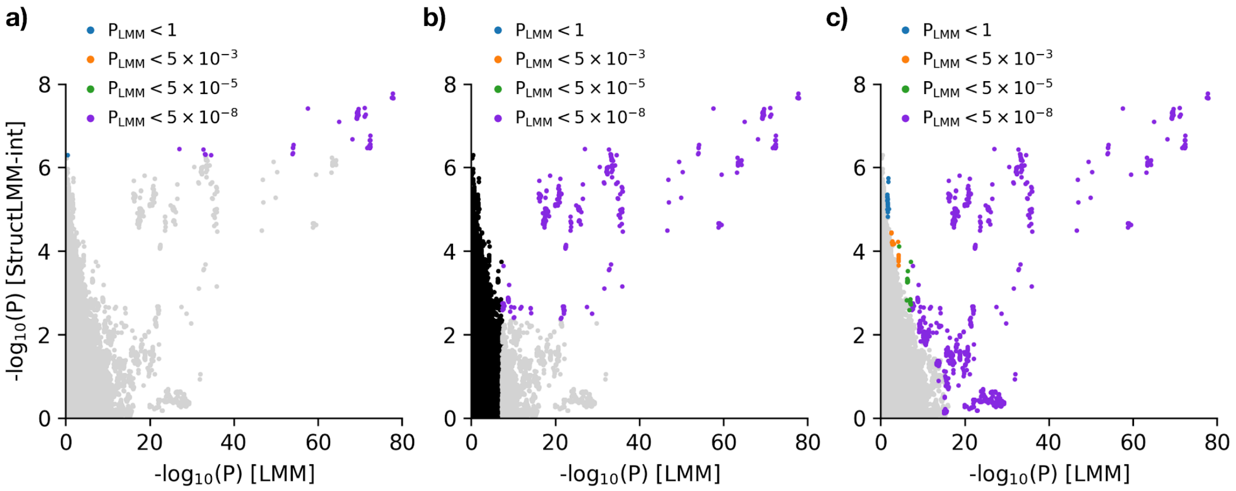


**Supplementary Figure 5 | Comparison of the variants identified with significant G×E interaction effects on the discovery set of UKBB individuals using alternative methods.** Scatter plots of genome-wide negative log StructLMM interaction P values (y-axis) versus negative log LMM association P values (x-axis, 7,515,856 variants), applied to UKBB BMI data using the discovery set of individuals (n = 126,077). Analogous to **Fig. 2** but variants with significant G×E interaction effects are coloured according to the association P value bin that they fall into. Considered were (**a**) results from a genome-wide G×E scan (7,515,856 variants), where significant interaction effects were defined using the 5% Benjamini-Hochberg false discovery rate, (**b**) results from conventional two-step filtering, where only variants (3,767 variants) with genome-wide significant ($P < 5 \times 10^{-8}$) association effects were taken forward for interaction testing (variants in black are those not tested for interaction effects) with significant interaction effects defined using the 5% Benjamini-Hochberg false discovery rate and (**c**) results from a genome-wide G×E scan (7,515,856 variants), where significant interaction effects were defined using the 5% conditional false discovery rate. Using the genome-wide FDR approach, the conventional two-step filtering approach and the cFDR approach resulted in 78, 330 and 964 variants corresponding to 3, 8 and 29 loci (LD clumped loci, $r^2 < 0.1$ within +/-500kb, **Methods**), respectively.

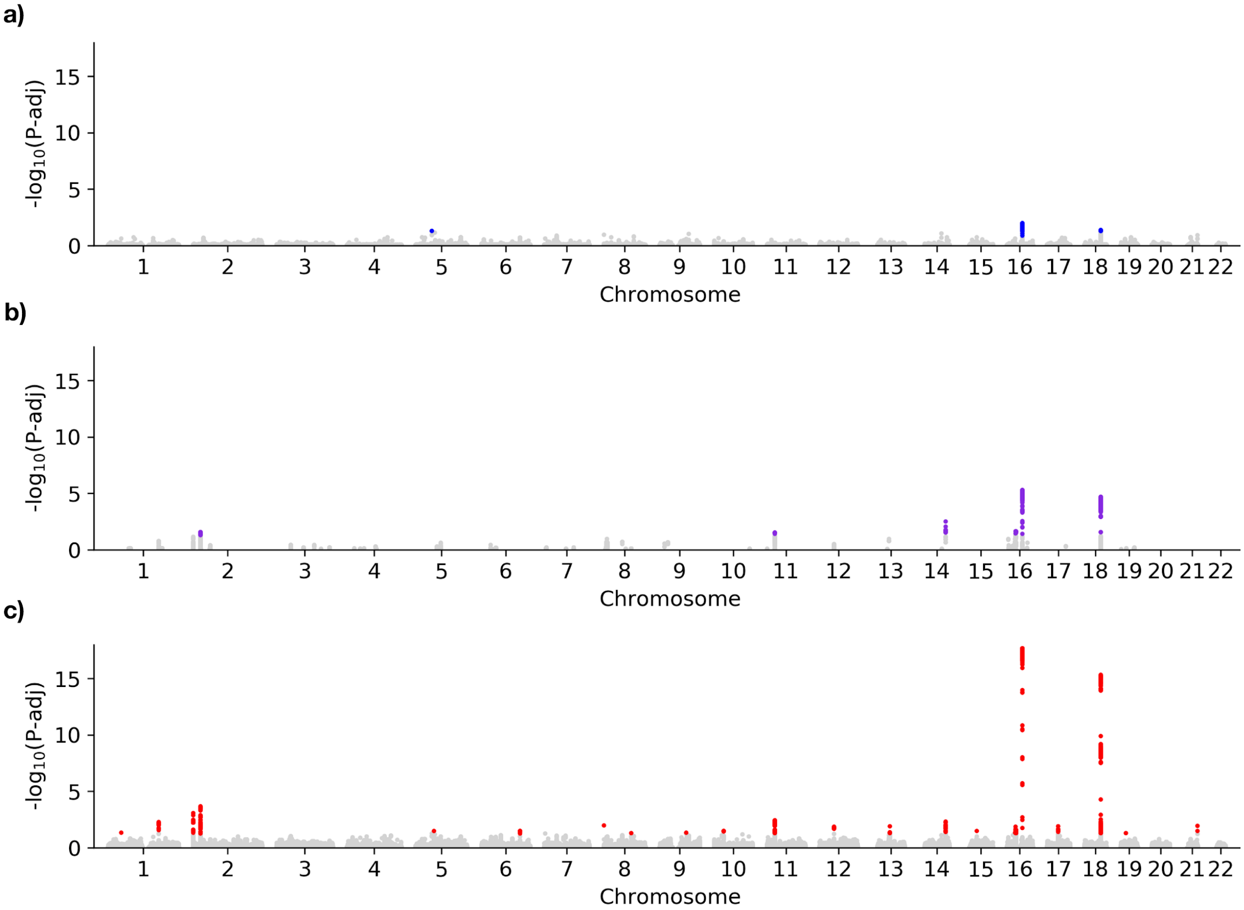


**Supplementary Figure 6 | Comparison of the variants and loci identified with significant G×E interaction effects using the discovery set of UKBB individuals for different methods.** Manhattan plots showing genome-wide negative log FDR adjusted P values (7,515,856 variants), applied to UKBB BMI data using the discovery set of individuals (n = 126,077). Considered were (**a**) results from a genome-wide G×E scan (7,515,856 variants), where significant interaction effects were defined using the 5% Benjamini-Hochberg false discovery rate, (**b**) results from conventional two-step filtering, where only variants (3,767 variants) with genome-wide significant ($P < 5 \times 10^{-8}$) association effects were taken forward for interaction testing with significant interaction effects defined using the 5% Benjamini-Hochberg false discovery rate and (**c**) results from a genome-wide G×E scan (7,515,856 variants), where significant interaction effects were defined using the 5% conditional false discovery rate. Using the genome-wide FDR approach, the conventional two-step filtering approach and the cFDR approach resulted in 78 (coloured in blue), 330 (coloured in purple) and 964 variants (coloured in red) corresponding to 3, 8 and 29 loci (LD clumped loci, $r^2 < 0.1$ within +/-500kb, **Methods**), respectively.

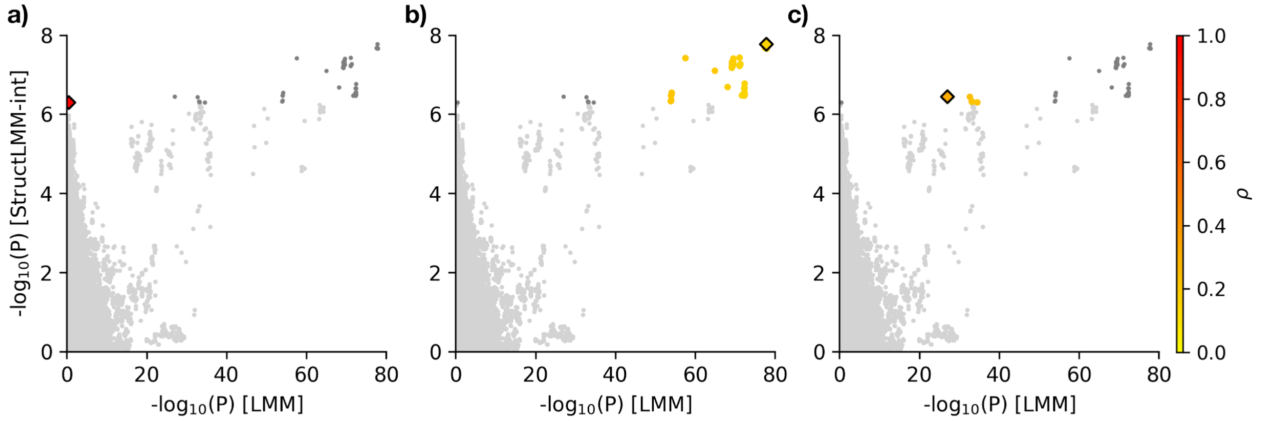


**Supplementary Figure 7 | Lead variants and corresponding variants in LD $r^2$ > 0.1 for loci identified with significant G×E interaction effects using the genome-wide FDR approach.** Scatter plots of genome-wide negative log StructLMM interaction P values (y-axis) against negative log LMM association P values (x-axis, 7,515,856 variants), applied to UKBB BMI data using the discovery set of individuals (n = 126,077), highlighting different loci. Variants within the focal loci (LD clumped loci, $r^2$ < 0.1 within +/-500kb, **Methods**) are coloured according to $\rho$, which estimates the fraction of the genetic variance due to G×E, with the lead variant per loci represented by a diamond. All other variants with significant interaction effects (Benjamini-Hochberg FDR < 5%) are displayed in dark grey. The three loci identified using the genome-wide FDR approach have lead variants at (**a**) 5:56865025, (**b**) 16:53806453 and (**c**) 18:57802714 (see **Supp. Table 1**).

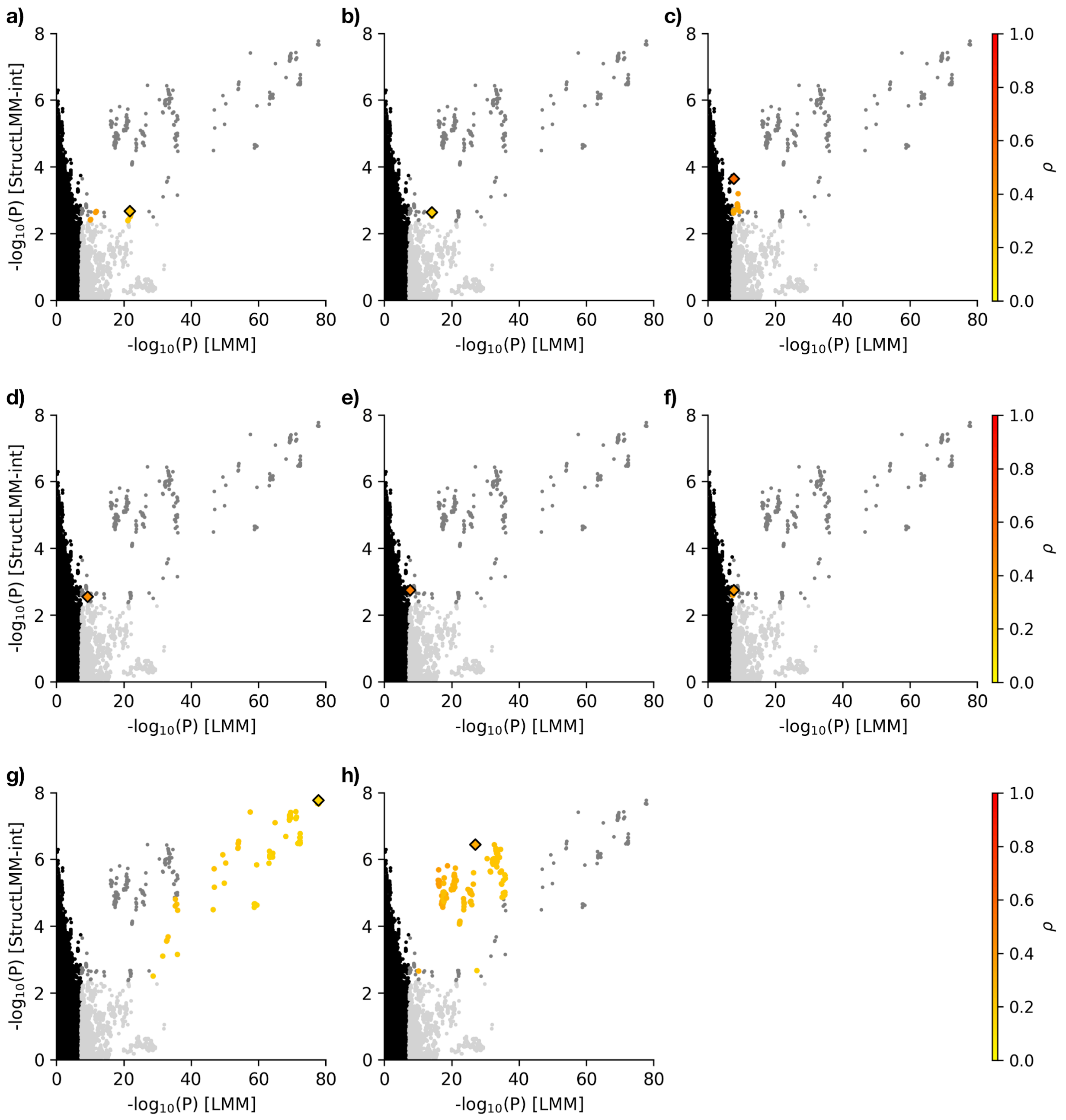


**Supplementary Figure 8 | Lead variants and corresponding variants in LD $r^2$ > 0.1 for loci identified with significant G×E interaction effects using the conventional two-step filtering approach.** Scatter plots of genome-wide negative log StructLMM interaction P values (y-axis) against negative log LMM association P values (x-axis, 7,515,856 variants), applied to UKBB BMI data using the discovery set of individuals (n = 126,077), highlighting different loci. Variants within the focal loci (LD clumped loci, $r^2$ < 0.1 within +/-500kb, **Methods**) are coloured according to $\rho$, which estimates the fraction of the genetic variance due to G×E, with the lead variant per loci represented by a diamond. All other variants with significant interaction effects (Benjamini-Hochberg FDR < 5%) are displayed in dark grey. The eight loci identified using the conventional two-step filtering FDR approach have lead variants at (**a**) 2:25158281, (**b**) 11:27723334, (**c**)

14:94120712, (**d**) 16:29001460, (**e**) 16:29984373, (**f**) 16:31347519, (**g**) 16:53806453 and (**h**) 18:57802714 (see **Supp. Table 1**).

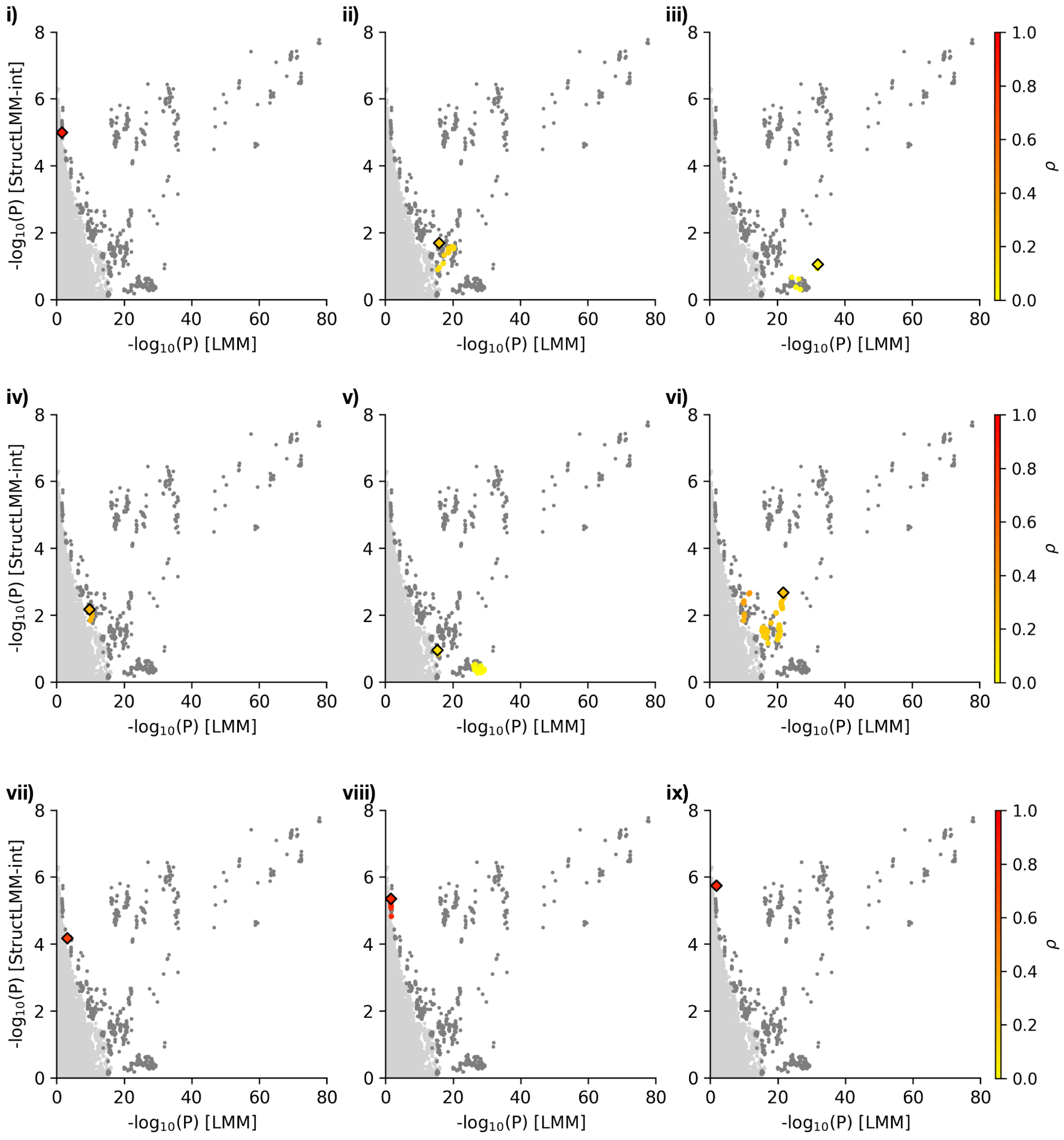

i)
ii)
iii)
iv)
v)
vi)
vii)
viii)
ix)
-log10(P) [StructLMM-int]
-log10(P) [LMM]
ρ

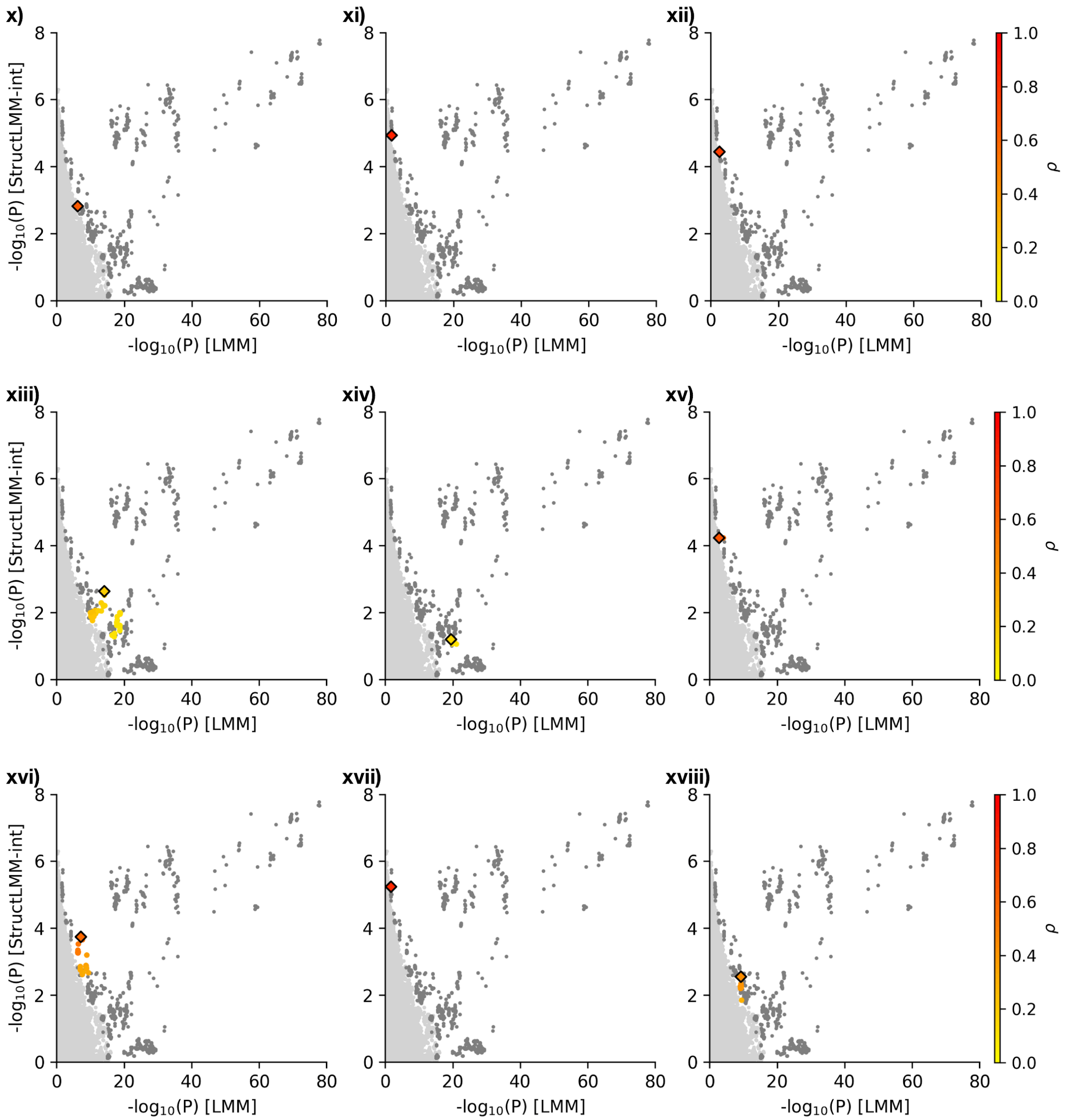
x)
xi)
xii)
xiii)
xiv)
xv)
xvi)
xvii)
xviii)
-log$_{10}$(P) [StructLMM-int]
-log$_{10}$(P) [LMM]
ρ

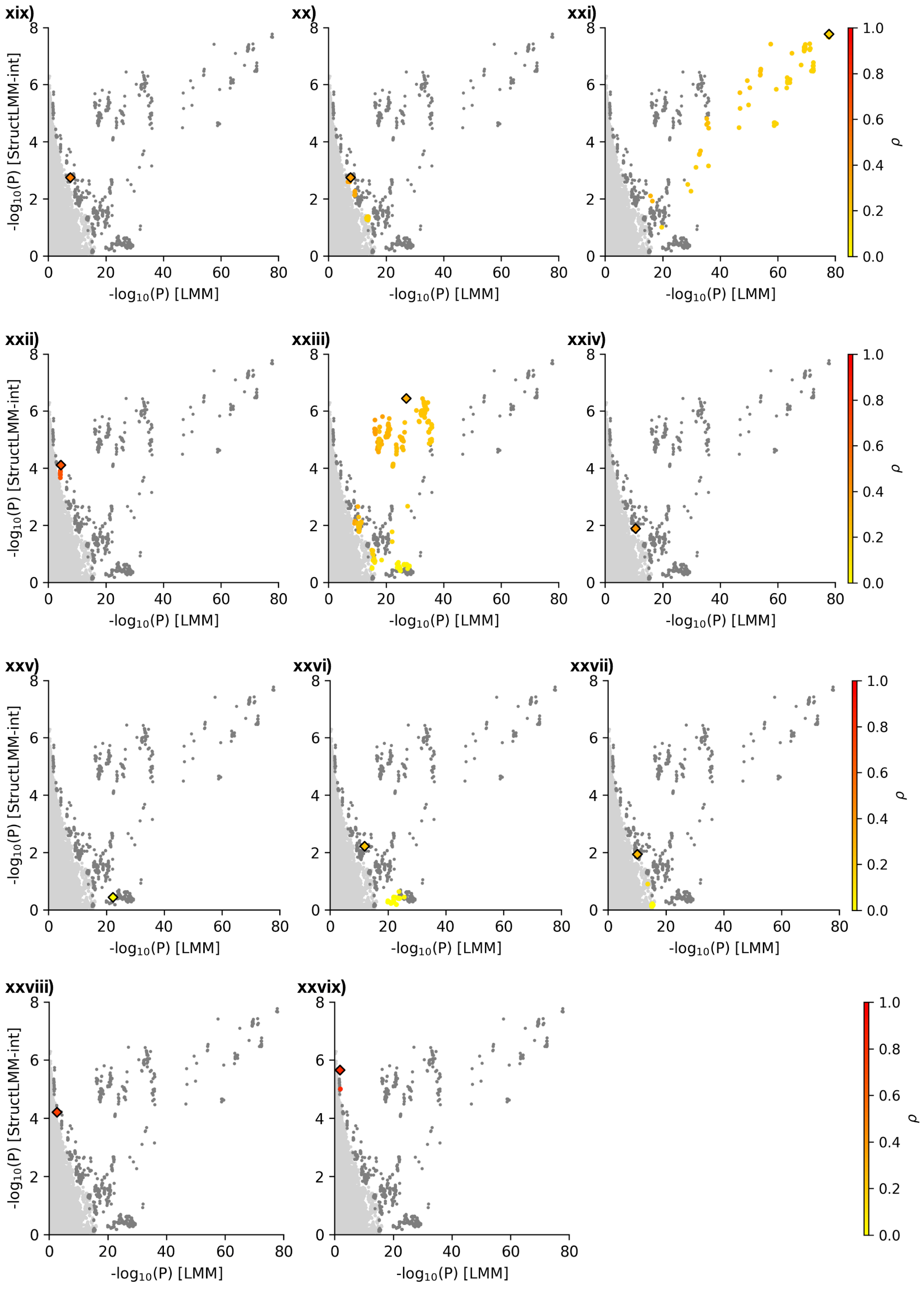

xix)
xx)
xxi)
xxii)
xxiii)
xxiv)
xxv)
xxvi)
xxvii)
xxviii)
xxvix)
-log10(P) [StructLMM-int]
-log10(P) [LMM]
0
20
40
60
80
0
2
4
6
8
ρ
0.0
0.2
0.4
0.6
0.8
1.0

**Supplementary Figure 9 | Lead variants and corresponding variants in LD $r^2 > 0.1$ for loci identified with significant G×E interaction effects using the cFDR approach.** Scatter plots of genome-wide negative log StructLMM interaction P values (y-axis) against negative log LMM association P values (x-axis, 7,515,856 variants), applied to UKBB BMI data using the discovery set of individuals (n = 126,077), highlighting different loci. Variants within the focal loci (LD clumped loci, $r^2 < 0.1$ within +/-500kb, **Methods**) are coloured according to $\rho$, which estimates the fraction of the genetic variance due to G×E, with the lead variant per loci represented by a diamond. All other variants with significant interaction effects (cFDR < 5%) are displayed in dark grey. The 29 loci identified using the cFDR approach have lead variants at (**i**) 1:46722389, (**ii**) 1:177887018, (**iii**) 2:417167, (**iv**) 2:433940, (**v**) 2:645190, (**vi**) 2:25158281, (**vii**) 5:63966889, (**viii**) 6:135918726, (**ix**) 8:203086, (**x**) 8:95628254, (**xi**) 9:92137422, (**xii**) 10:33299356, (**xiii**) 11:27723334, (**xiv**) 12:50246252, (**xv**) 13:62619925, (**xvi**) 14:94132712, (**xvii**) 15:45093429, (**xviii**) 16:29001460, (**xix**) 16:29984373, (**xx**) 16:31347519, (**xxi**) 16:53806453, (**xxii**) 17:39262850, (**xxiii**) 18:57802714 (**xxiv**) 18:57913703, (**xxv**) 18:57955945, (**xxvi**) 18:57967655, (**xxvii**) 18:58049656, (**xxviii**) 19:18198063, (**xxvix**) 21:46695006 (see **Supp. Table 1**).

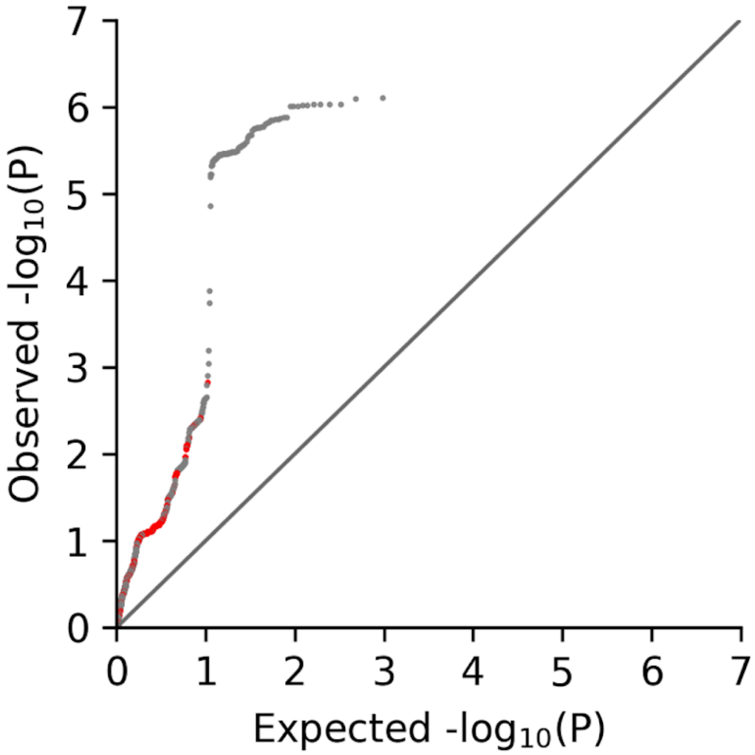


**Supplementary Figure 10 | QQ plot of the StructLMM interaction test for the subset of variants identified in discovery using the validation dataset.** QQ plot of negative log P values from the StructLMM interaction test, using individuals in the validation dataset (n = 126,076) for the 964 variants identified by the cFDR approach in the discovery analysis. Analogous to **Fig. 3a** but all variants corresponding to the eight loci (LD clumped loci, $r^2 < 0.1$ within +/-500kb, **Methods**) also identified by the conventional two-step filtering and/or the genome-wide FDR approach in discovery are displayed in grey whilst variants corresponding to the 21 loci exclusively identified by the cFDR approach in discovery are displayed in red.

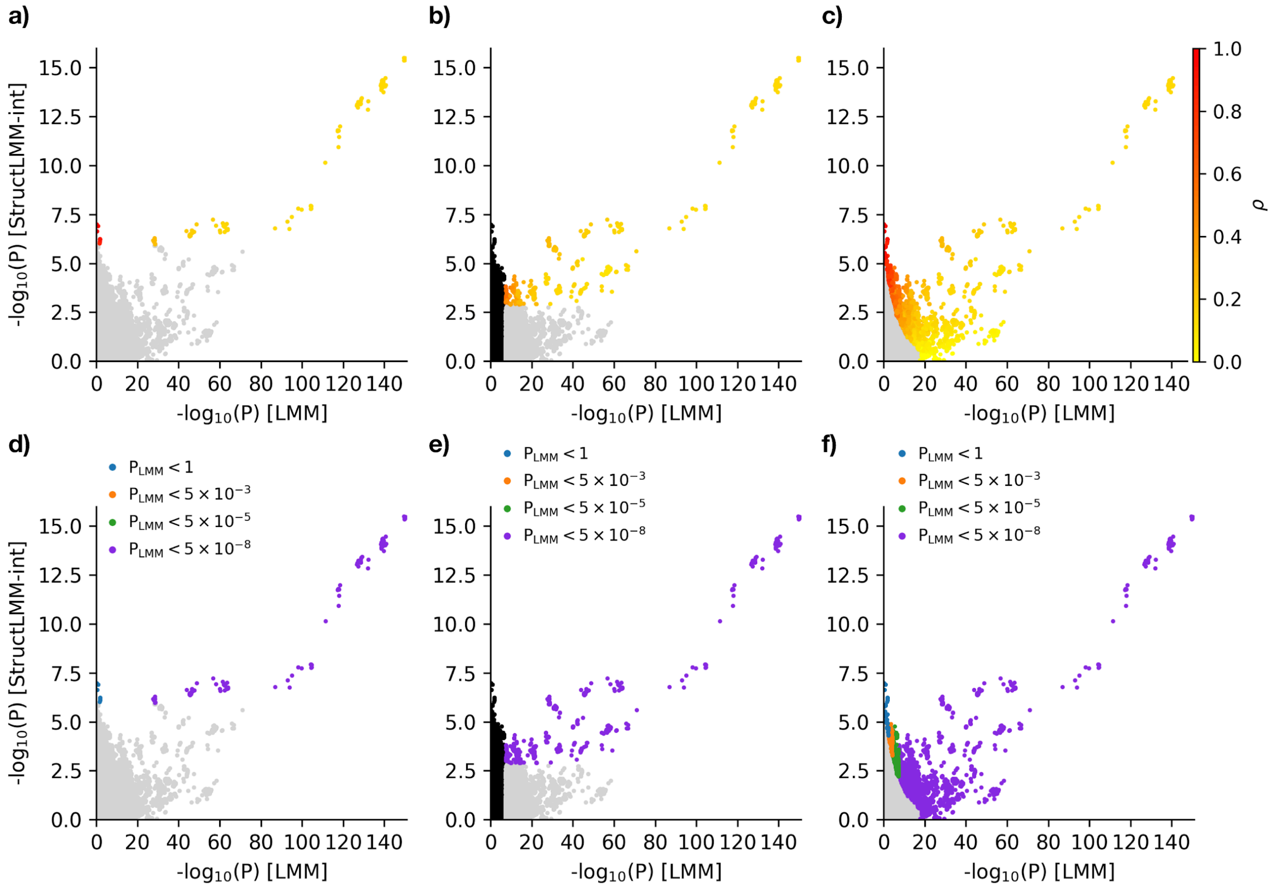


**Supplementary Figure 11 | Comparison of the variants identified with significant G×E interaction effects using the full set of UKBB individuals for different methods.** Scatter plots of genome-wide negative log StructLMM interaction P values (y-axis) against negative log LMM association P values (x-axis, 7,515,856 variants), applied to UKBB BMI data using the full set of individuals (n = 252,188). Variants with significant G×E interaction effects are coloured according to (**a-c**) $\rho$, which estimates the fraction of the genetic variance due to G×E and (**d-f**) the association P value bin that they fall into. Considered were (**a**, **d**) a genome-wide approach where all variants (7,515,856 variants) were tested for interaction effects with significant interaction effects defined using the 5% Benjamini-Hochberg false discovery rate, (**b**, **e**) conventional two-step filtering approach where only variants (17,606 variants) with genome-wide significant ($P < 5 \times 10^{-8}$) association effects were taken forward for interaction testing (variants in black are those not tested for interaction effects) with significant interaction effects defined using the 5% Benjamini-Hochberg false discovery rate and (**c**, **f**) a genome-wide approach where all variants (7,515,856 variants) were tested for interaction effects with significant interaction effects defined using the 5% conditional false discovery rate. Using the genome-wide FDR approach, the conventional two-step filtering approach, and the cFDR approach resulted in 164, 451 and 4,179 variants corresponding to 6, 23 and 140 loci (LD clumped loci, $r^2 < 0.1$ within +/-500kb, **Methods**), respectively.

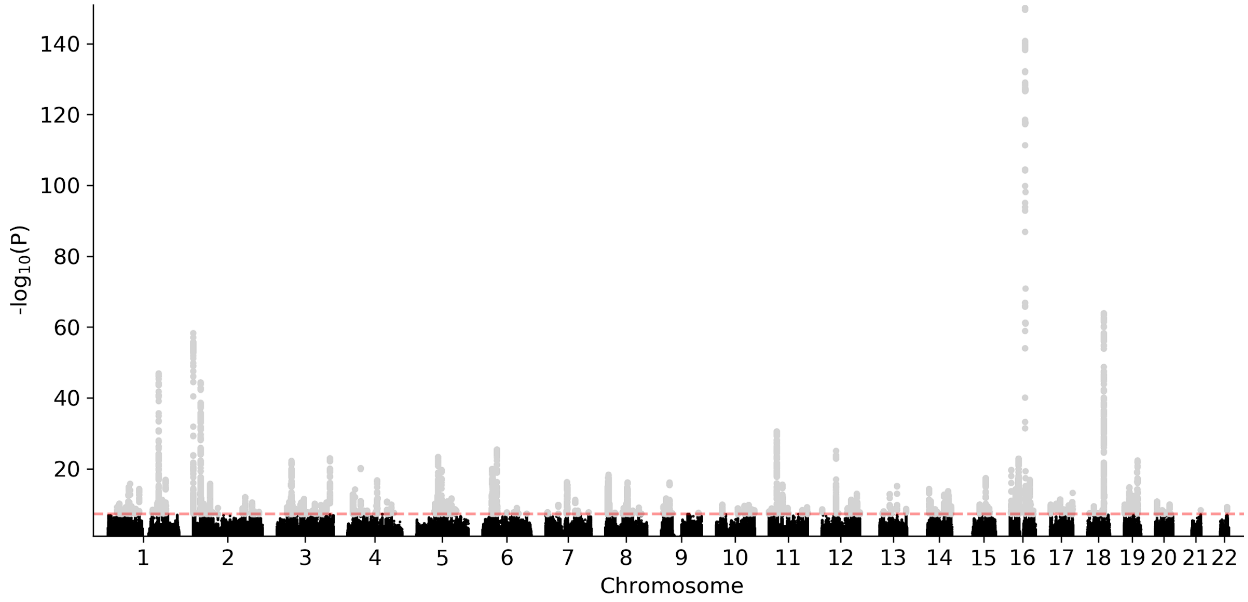


**Supplementary Figure 12 | Association test results applied to the full set of UKBB individuals.** Manhattan plot showing genome-wide negative log P values (7,515,856 variants) obtained from the association test, LMM, applied to UKBB BMI data using the full set of individuals (n = 252,188). The dashed red line denotes the genome-wide significance threshold ($P < 5 \times 10^{-8}$). Genome-wide significant variants (grey) are those tested for interaction testing using the two-step filtering approach. 17,606 variants, corresponding to 327 loci (LD clumped loci, $r^2 < 0.1$ within +/-500kb, **Methods**) are genome-wide significant.

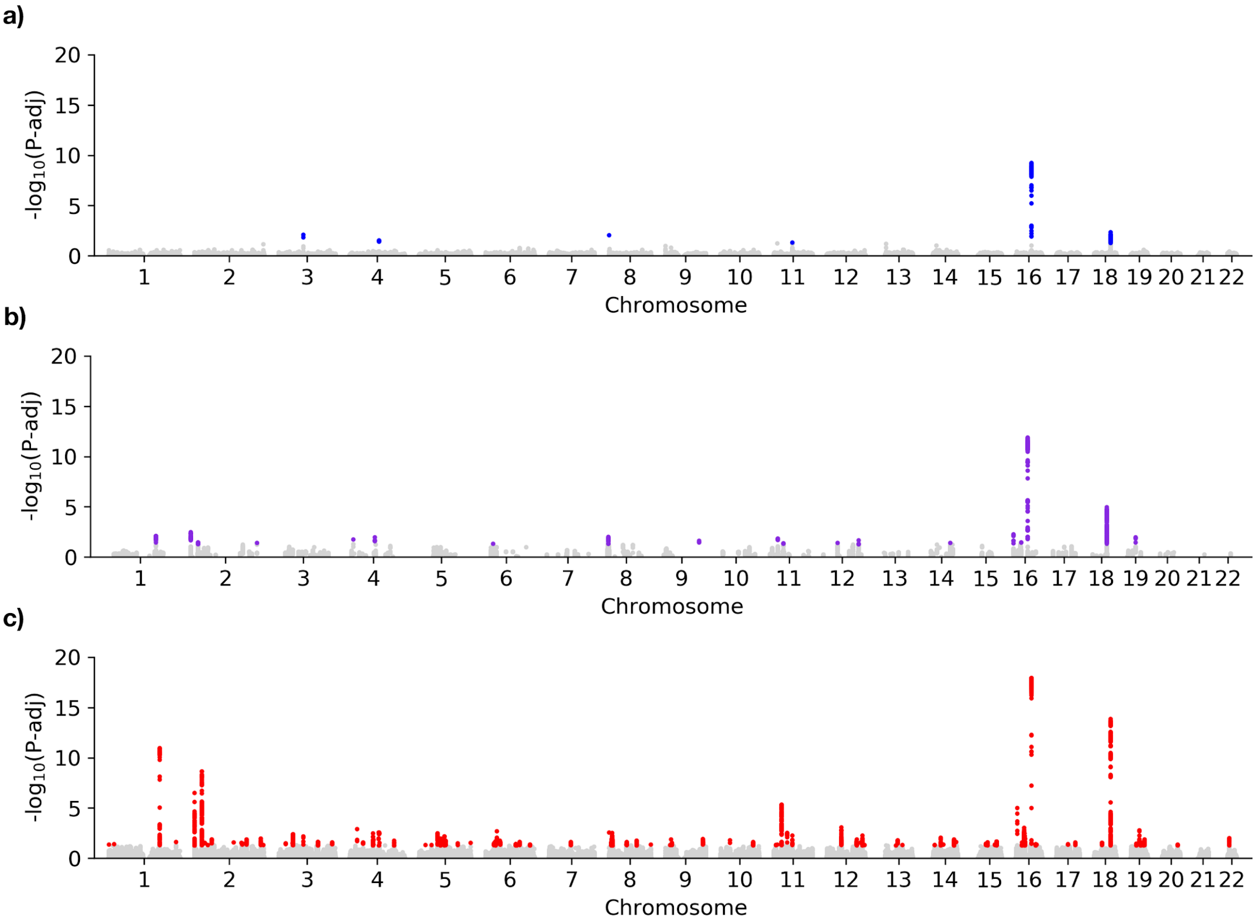


**Supplementary Figure 13 | Comparison of the variants and loci identified with significant G×E interaction effects using the full set of UKBB individuals for different methods.** Manhattan plots showing genome-wide negative log FDR adjusted P values (7,515,856 variants), applied to UKBB BMI data using the full set of individuals (n = 252,188). Considered were (**a**) a genome-wide approach where all variants (7,515,856 variants) were tested for interaction effects with significant interaction effects defined using the 5% Benjamini-Hochberg false discovery rate, (**b**) a conventional two-step filtering approach where only variants (17,606 variants) with genome-wide significant ($P < 5 \times 10^{-8}$) association effects were taken forward for interaction testing with significant interaction effects defined using the 5% Benjamini-Hochberg false discovery rate and (**c**) a genome-wide approach where all variants (7,515,856 variants) were tested for interaction effects with significant interaction effects defined using the 5% conditional false discovery rate. Using the genome-wide FDR approach, the conventional two-step filtering approach and the cFDR approach resulted in 164 (coloured in blue), 451 (coloured in purple) and 4,179 variants (coloured in red) corresponding to 6, 23 and 140 loci (LD clumped loci, $r^2 < 0.1$ within +/-500kb, **Methods**), respectively.

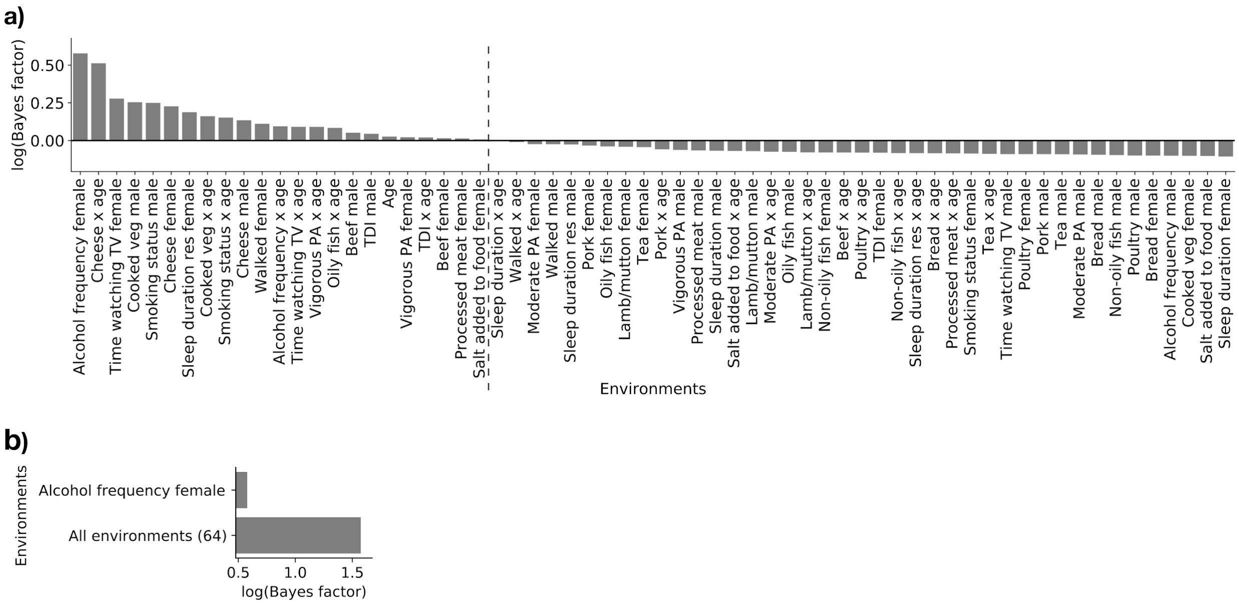


**Supplementary Figure 14 | Exploration of the putative driving environments of the G×E effect at the *FAIM2* loci using the validation lead variant (rs7132908) and validation set of UKBB individuals.** Relevance of individual environmental variables for the G×E effect at the validation lead SNP, rs7132908 (*FAIM2* loci), showing Bayes factors between the model that contains all 64 environments and models with environmental variables removed using the set of UKBB validation individuals (n = 126,076). Shown are (**a**) results based on removing single environmental variables ordered by Bayes factor, with those environments with positive evidence of driving the G×E displayed to the left of the dashed line, whilst those with negative evidence are to the right (**Methods**) and (**b**) the result from using a (greedy) backward elimination stopping when there is evidence that the selected environments explain the observed G×E effect (**Methods**); for comparison, shown is the total evidence of all 64 environmental variables. At this SNP there is evidence that alcohol intake frequency in women is sufficient to explain the observed G×E effect.
